# What kind of birds are more susceptible to avian malaria? A global analysis based on interpretable machine learning approach

**DOI:** 10.1101/2025.11.02.686156

**Authors:** Xinyi Wang, Swapnil Mishra, Lu Dong

## Abstract

**Aim:** Avian malaria (genus *Plasmodium*) is a mosquito-transmitted parasitic disease of birds. It has a wide distribution across the world, infecting more than 2,400 bird species, posing a great threat to bird health. However, over half of bird species have a cumulative sample size of fewer than 20 individuals, leading to a limited understanding of the global patterns and mechanisms of their susceptibility to avian malaria. Our aim is to use interpretable machine learning to identify the global ecological and evolutionary drivers shaping species-level avian malaria susceptibility in birds.

Location Global.

Time period 1994–2023.

Major taxa studied

Global bird species and their malaria parasites (genus *Plasmodium*)

**Methods:** Based on infection data and traits from 72,406 birds of 2,544 species worldwide, we developed an interpretable machine learning (IML) approach to identify the global drivers of species-level susceptibility and their trends. We further applied our model to predict the susceptibility of bird species with a sample size of less than 30 and tested multiple hypotheses related to differences in parasite susceptibility in birds.

**Results:** Our model distinguished susceptible birds with moderate accuracy (F1 score = 0.72) and predicted 752 bird species to be highly susceptible to avian malaria, including 16 threatened species. Susceptibility showed a moderate but significant phylogenetic signal, with most susceptible species belonging to Passeriformes. Highly susceptible species were generally characterized by larger body size, omnivory, ground-foraging behavior, wider geographic ranges, and medium diversification rates.

**Main Conclusions:** We show that ecological and evolutionary factors together shape species-level susceptibility of birds to avian malaria. Interpretable machine learning integrating host traits offers a global insights into infection mechanisms and supports the development of more precise surveillance and control strategies.

## 1 INTRODUCTION

Pathogens present significant challenges to wildlife biodiversity [82, 89]. Understanding host susceptibility is essential for predicting and mitigating these impacts, as well as for assessing the risk of pathogen spillover from wildlife to domestic animals and humans [23]. Susceptibility, broadly defined as ‘proneness to infection’, is a valuable metric for describing inter-specific differences in the host’s ability to respond to pathogen infections [5, 30, 70, 79]. Therefore, a major goal in disease ecology is to find out inter-specific susceptibility patterns and their drivers among hosts for explaining the emergence and evolution of infectious diseases, particularly in the context of global environmental changes [7].

Host susceptibility is determined by the epidemiological characteristics, including exposure (e.g., direct contact, environmental uptake, and vector contact) and host competence (i.e., a host’s ability to acquire and transmit pathogens) [17, 42]. Here, we use the prevalence as the index of susceptibility [6], which reflects both the exposure and host competence process of transmission. Multiple hypotheses have been proposed to explain the correlation between host ecological traits and inter-specific variation in susceptibility to pathogens [97]. These hypotheses have been proposed to explain their correlations in various aspects, including dispersal capability, habitat use, and life history (Table 1). For vector-borne diseases, habitat use is closely linked to exposure levels [61, 73, 76]. For example, animals foraging on the understory strata are more likely to be infected by malaria due to the feeding preferences of mosquitoes [11, 29, 63]. Life-history tradeoffs and parasite adaptation can influence host competence [50]. The life-history theory suggests trade-offs between investment in reproduction and immune defense against infection [84]. Fast-living species, which are characterized by small body size, rapid reproduce, and short lifespans, may be more susceptible to pathogens [2, 90, 101]. Such relationships have been documented in various zoonotic infectious diseases, including avian influenza [101], Lyme disease (*Borrelia burgdorferi*), West Nile Virus [41], etc. Migration, as one of the most important strategies to maximize individual fitness [3], is related to both exposure and host competence. Migratory hosts can span broad-scale geographical regions, which can either enhance the geographic spread of many pathogens or avoid infected habitats [3, 56, 78]. Also, the energy costs associated with migration can reduce immunity, thus potentially leading to the removal of infected individuals from the population [10].

**Table 1.**
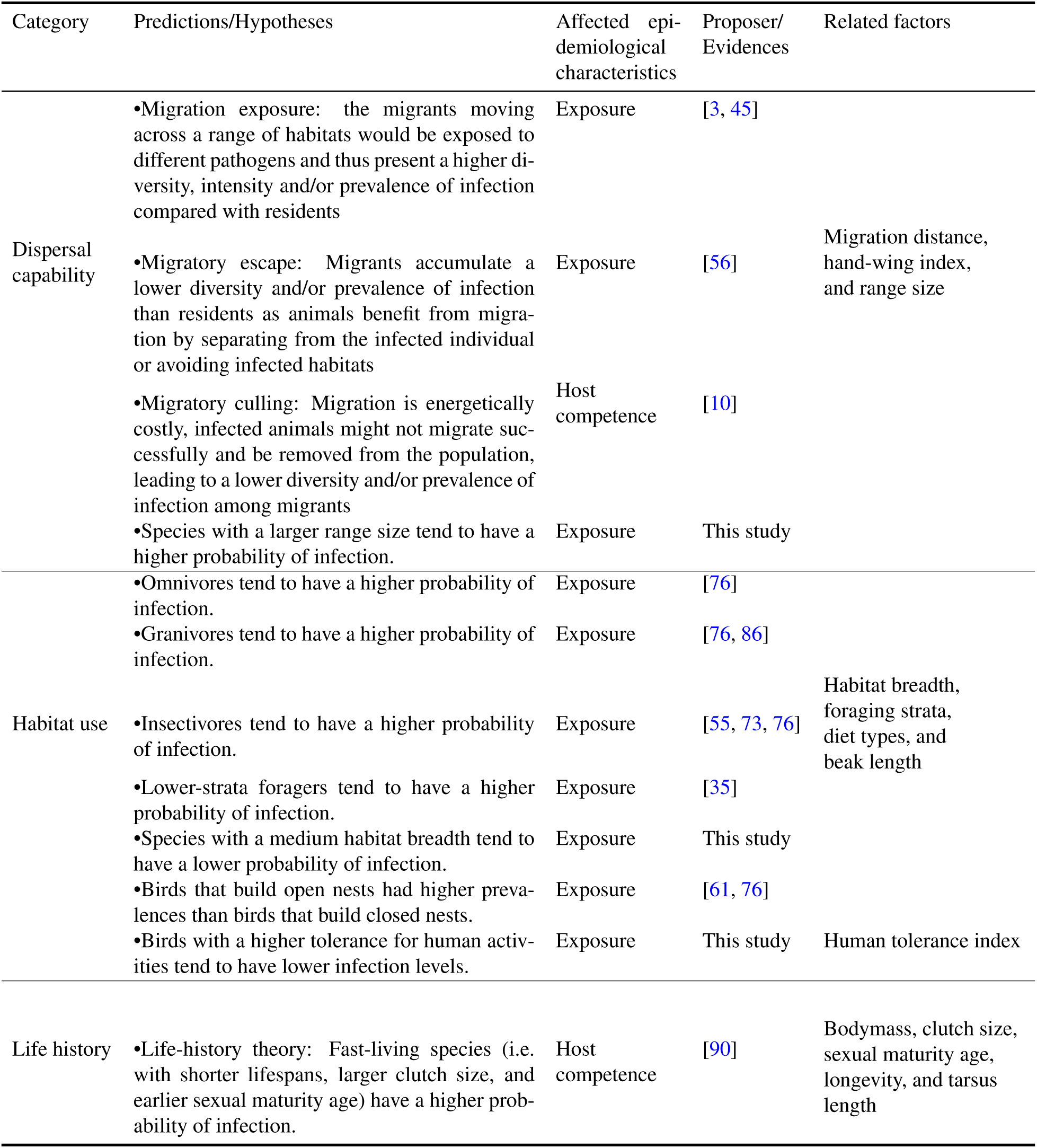
Ecological factors and related hypotheses that influence inter-species susceptibility in birds.

The evolutionary relationship between hosts and parasites adds further complexity to susceptibility [6]. First, the ‘lability hypothesis’ suggests that susceptibility is labile across the extent of multi-host, multi-parasite systems [6], given that significant short-term and spatial variation in both exposure to pathogens and host immune defensive abilities [85, 99]. This hypothesis predicts a lack of phylogenetic signal. Second, host evolutionary processes may affect host susceptibility to pathogens. Host susceptibility has shown a conserved pattern of evolution across the host phylogeny in some systems (e.g. plant-mistletoe [102]; amphibian-fungal pathogen [33]), possibly due to the phylogenetically conserved host immune systems and traits affecting interspecific variation in susceptibility [6, 20, 40, 57, 65]. Faster-diversifying clades may have higher prevalence because host shifts tend to happen among close relatives [21]. Under the second hypothesis, we would expect susceptibility to be phylogenetically conserved and influenced by host diversification rates. Third, parasites, in turn, may act as a strong selective pressure in shaping key aspects of host evolutionary history, including speciation [52, 69]. The Red Queen hypothesis further emphasizes the dynamic nature of host-pathogen interactions, describing cycles of adaptation and counteradaptation that shape both host and pathogen evolutionary histories. [92].

It is widely recognized that incorporating multi-host, multi-parasite interactions is crucial for better understanding and predicting disease threats [25]. With the data-sharing efforts of the research community, vast amounts of data have been generated and accessible through public databases such as GISAID and MalAvi [8, 81]. However, disease ecology studies remain challenging due to the intricate interplay of factors across various biological scales and the interwoven evolutionary relationships between hosts and pathogens [42]. On one hand, statistical models have limitations as they rely on assumptions, such as low dimensionality of input data, which may fail to hold when applied to complex real-world datasets. On the other hand, machine learning (ML) algorithms, capable of processing vast amounts of data from diverse sources with greater predictive power, often lack transparency and interpretability [43]. Interpretable machine learning (IML) emerges as a promising approach for exploring complex systems, combining the predictive strength of ML with feature-level interpretability [39, 51, 66]. IML has been successfully applied in various fields such as disease prediction [26], genomic landscape analysis [25], and biomarker identification [64]. In this study, we developed an IML framework that integrates ML models and a Shapley value-based component [80].

Avian malaria is a globally distributed mosquito-borne disease caused by *Plasmodium* parasites. It serves as a classic multi-host, multi-pathogen system for investigating host-pathogen interactions, coevolutionary processes, and the role of pathogens in host life-history evolution [37, 54, 74]. Infecting over 2,400 bird species [15, 91], [24], avian malaria has caused significant impacts on bird populations, including the endangerment and extinction of several Hawaiian songbird species in the first half of the 20th century [93, 94]. Moreover, chronic infections can significantly decrease the fitness of birds through telomere degradation and senescence [4]. The complexity of the avian malaria system, involving diverse vectors, hosts, and environments, makes it an ideal model for studying host susceptibility patterns. [54] [91]. Some studies have explored the relationship between susceptibility and one or a few of the above factors [6, 62, 71, 100], but due to limitations in data accumulation and analytical methods, no previous analysis has comprehensively examined the complex links between macroscopic traits, and evolutionary process and species-level host susceptibility in the global scale.

In this study, we compiled a comprehensive global dataset of 72,406 birds, together with their malaria infection status and host traits across functional, biogeographical, and evolutionary dimensions. By developing a methodological framework based on state-of-the-art IML methods, we aim to (1) identify the key ecological and evolutionary factors shaping malaria susceptibility in birds, and test related hypotheses;(2) predict the susceptibility of globally under-sampled bird species, particularly threatened species, thereby providing scientific guidance for avian biodiversity conservation and pathogen surveillance.

## 2 MATERIALS AND METHODS

### 2.1 Avian malaria infection data

To collect global infection data of avian *Plasmodium* parasites, we compiled data from three sources. First, we utilised a bird infection dataset based on the Malavi Database [8], as compiled by [24]. This global dataset contains presence-absence records from 53,669 birds from 1994 to 2019. Second, as the above dataset doesn’t contain data from 2021 onwards, we retrieved records of avian *Plasmodium* infection records from the MalAvi database (Version 2.5.8, accessed on Dec 20, 2023) from 2021 to 2023 [8]. We filtered out records without sampling or infection numbers. Records with infected counts exceeding the sampling counts were deleted, indicating a possible error. We only considered wild birds, so captive individuals (n=1,449) and nestlings (n=243) were also deleted. What’s more, we enriched the data with new samples collected by our group, which ensures the representativeness of birds from the East Asia region. We added 1,222 records collected by our research group in northern China between 2014 and 2021. Subsequently, the overall prevalence of each species was calculated based on data synthesized from the aforementioned three components.

### 2.2 Avian trait data

We compiled a dataset of avian hosts that integrates both ecological and evolutionary traits associated with susceptibility. Body mass, beak length, tarsus length, tail length, wing length, and range size were extracted from the AVONET databank [88]. The diet data was derived from EltonTraits v.1.0 [96], which details the percentage utilization of seven diet types in each species’ diet: (1) invertebrates, (2) vertebrates, (3) fish, (4) fruit, (5) nectar, (6) seed and (7) other plant material. The foraging strata was characterized by the percentage of time spent on four types of habitats: (1) foraging above the canopy, (2) foraging in mid to high levels in trees, (3) foraging below 2m in the understory, and (4) foraging on ground [96]. For behavior traits, the Hand-wing index (HWI) was obtained from the AVONET databank [88]. HWI is a morphological metric linked to wing aspect ratio and is widely used as a single-parameter proxy of avian flight efficiency and dispersal ability. High HWI values indicate elongated wings and better flight performance, while low HWI values are associated with shorter, broader wings and inferior flight performance. Migration distance was collected from [19]. For resident species, this value was set to zero. Habitat breadth data were sourced from [18], reflecting the degree of habitat specialization by quantifying species co-occurrences across 101 habitat categories described by the International Union for the Conservation of Nature (IUCN). We also included the Human Tolerance Index (HTI) to measure bird species’ tolerance to human pressures, which may also affect parasitic infections [60].

The evolutionary traits of hosts included diversification rate (DR) and phylogenetic category. DR measures how quickly new species are formed in a lineage. Species with higher DR have shorter and more recent branches shared with other species, indicating a fast speciation rate. Conversely, species with lower DR have longer and more unique branches [103]. We randomly created a sample of 1000 phylogenetic trees of global birds (9993 species; the ‘Hackett’ backbone; [46]) from VertLife (http://vertlife.org/phylosubsets) [103], and then calculated the mean DR using the ’epm’ R package(version 1.1.2) [87]. We also classified the genome-scale phylogenic tree [44] into 13 main categories (Figure S1) to represent avian taxonomy.

We also queried the global population trends and IUCN red list categories of threat levels from IUCN Red List (v2024-1) (https://www.iucnredlist.org/). Global population trends encompassed increasing, stable, and decreasing categories. In the IUCN Red List, Least-Concern species are considered not threatened (0), while those ranked above Least Concern (i.e. Near Threatened, Vulnerable, Endangered, and Critically Endangered) are defined as threatened (1).

We tested all traits for collinearity, retaining the more ecologically integrative variable when correlations exceeded 80%. As tail length and wing length were highly correlated with the handwing index, which is widely used to measure avian flight ability, only the hand-wing index was retained. In total, 23 host traits were included in the machine learning model.

### 2.3 Model development

In this study, we constructed a framework based on interpretable machine learning to predict avian susceptibility to malaria using host traits (Figure 1). Bird species with higher prevalence were considered to be more susceptible to avian malaria. To ensure robustness, we trained models on well-sampled birds and then applied them to under-sampled birds.

**Fig 1.**
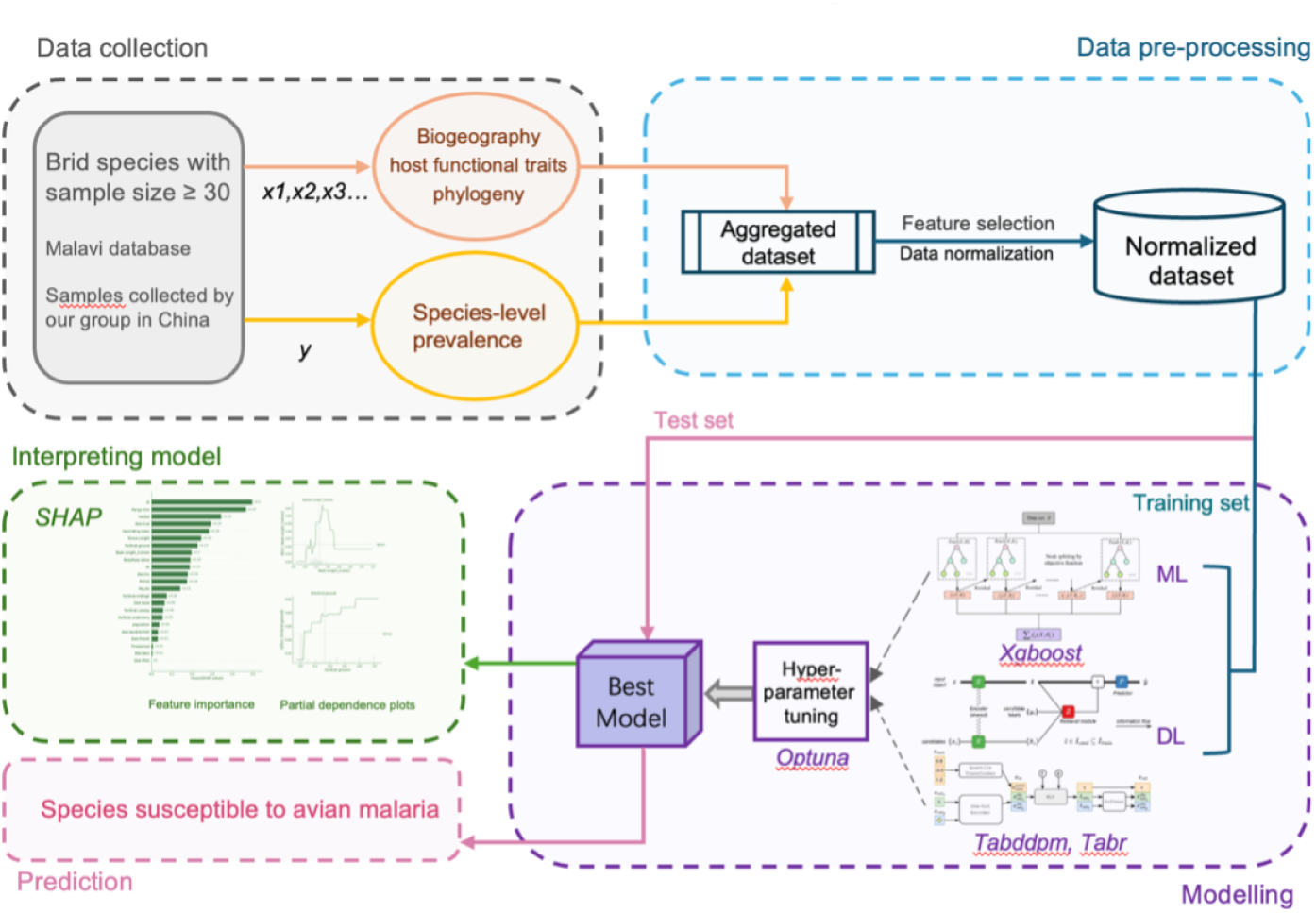
Flow chart showing the main steps in our IML framework.

We applied three ML/DL models that are either widely recognized or recently developed: eXtreme Gradient Boosting (XGBoost) [13], TabR [32], and TabDDPM [53]. XGBoost is a well-established scalable end-to-end tree boosting system known for its performance and efficiency [13], making it widely recognized as the algorithm of choice for many winning teams of machine learning competitions. TabR and TabDDPM represent the latest advancements in deep learning models specifically designed for handling tabular data problems [32, 53]. TabR incorporates a feed-forward network structure, featuring a custom component similar to k-Nearest Neighbors in its core [32]. TabDDPM adapts the Denoising Diffusion Probabilistic Model (DDPM) framework [38, 83], traditionally used for generative tasks, to address tabular data challenges [53].

Bird species with a sample size exceeding 30 were reasonably defined as well-sampled (58,251 bird individuals, covering 457 species, accounting for 80.4% of the total sampling records). Detailed sensitivity analysis was presented in Table S1 and Supplementary methods. We randomly split our data into a training set (80%) and a validation set (20%). We used K-fold cross-validation with the 80/20 split for hyperparameter optimization and performance [67]. The automated scripts of TabR and TabDDPM come with built-in hyperparameter tuning functionality. For XGBoost, we utilized the Optuna software [1] to do the automated hyperparameter tuning. We chose the TreeStructure Parzen Estimator (TPE), a Bayesian optimization (BO) algorithm configured in Optuna to sample hyperparameters [9]. We used the F1 score to gauge the capability of models because F1 score provides a balance between precision (i.e. true positives/(true positives + false positives)) and recall (sensitivity, i.e. true positives/(true positives + false negatives)). Two rounds of optimization were conducted to maximize the F1 score, each comprising 1000 trials. The detailed hyperparameters and their sample space are shown in Table S2 and Figure S2. For TabR and TabDDPM, their automated scripts come with built-in hyperparameter tuning.

We applied the best model to the full trait dataset of under-sampled avian species to identify those susceptible to avian malaria. To assess the phylogenetic signal in predictions, we calculated Pagel’s *λ* of probabilities using the adephylo R package (v.1.1-16) [49]. The predicted susceptibility was mapped onto the phylogenetic tree as a trait using phytools R package (v.2.1.1) [72].

Finally, SHAP (Shapley Additive Explanations) python package was used to identify important features by calculating the Shapley values [80] of the best model [58]. SHAP is a game theoretic approach widely used in explaining how machine learning models apply to predictions [12, 31]. This approach links optimal credit distribution to localized explanations by employing the Shapley values from game theory and their associated extensions [58]. We illustrated trait profiles through the use of SHAP partial dependence plots. These plots delineate how each feature’s contribution to the model’s predictions varies across its values.

## 3 RESULTS

### 3.1 Global sampling and infection status of avian malaria parasites

72,406 wild bird individuals were included in the final dataset, of which 10,359 tested positive for *Plasmodium* parasites. The estimated empirical global overall prevalence was 14.34% (CI: 14.06% –14.8%). 2,539 bird species have tested for avian *Plasmodium*, among which 1,118 species(44.0%) were found to have at least one record of avian *Plasmodium* infection. The average number of samples per bird species was 28.6, with a median of 6, and quartiles of 19 and 2. Some top well-sampled species included Silvereye (*Zosterops lateralis*), Grey Catbird (*Dumetella carolinensis*), Austral Thrush (*Turdus falcklandii*), and House Sparrow (*Passer domesticus*) (Figure 2). Some species with high prevalence included Morepork (*Ninox novaeseelandiae*, 100%, n = 36), Song Sparrow (*Melospiza melodia*, 60%, n = 420), and Godlewski’s bunting (*Emberiza godlewskii*, 78%, n = 245) (Figure 2).

**Fig 2.**
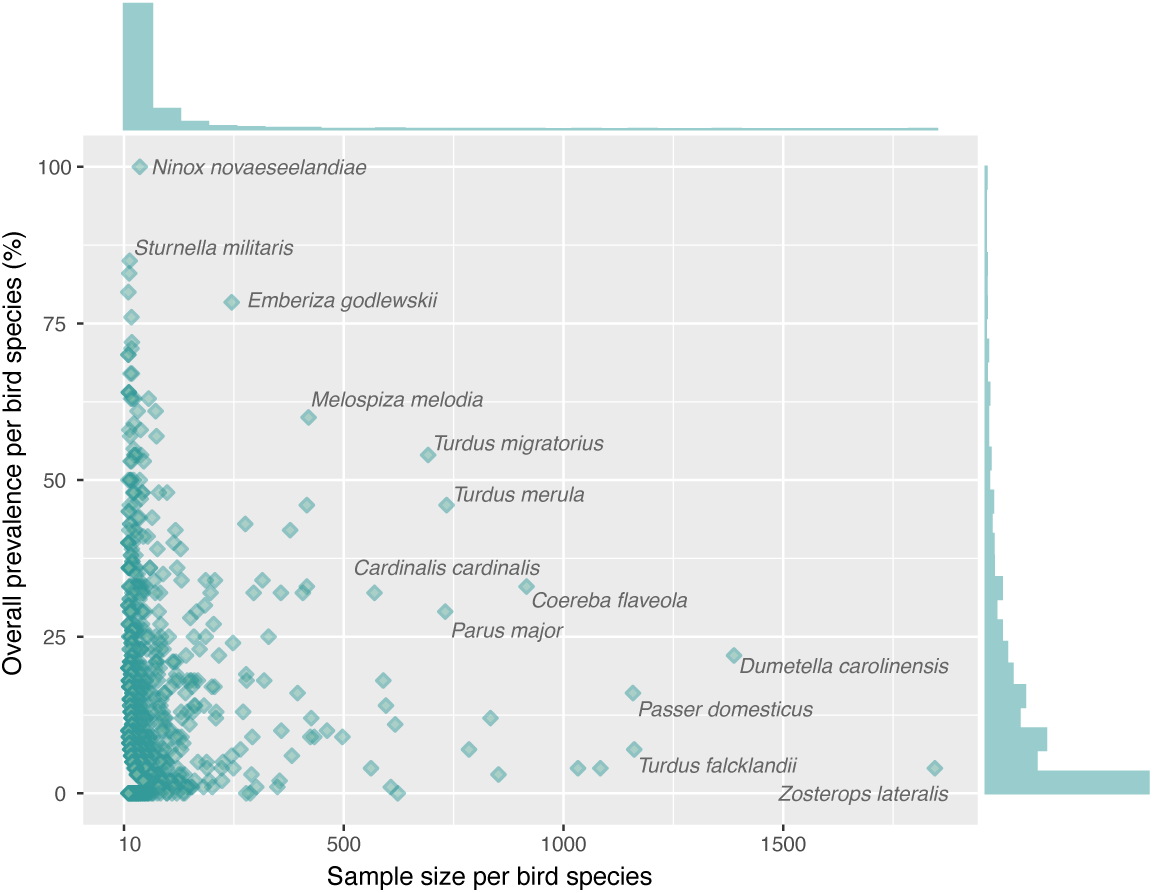
Plot showing the global sampling size of each bird species and its infection status of avian malaria parasites. Species with high prevalence and/or large sampling volumes are labeled.

### 3.2 Bird species highly susceptible to avian malaria

Based on the interpretable machine learning model, a total of 752 bird species were identified as highly susceptible (Supplementary Table 2), accounting for 36.4% of all tested species. Highly susceptible taxa were mainly concentrated within Passeriformes (Figure 3), including Ploceoidea, Muscicapidae, Thraupidae, Emberizidae, Passerellidae, Icteridae, Dicaeidae, Motacillidae, Fringillidae, Pycnonotidae, Passeridae, Corvidae, and Laniidae (Figure 3). Among non-passerines, families with elevated susceptibility included Accipitridae, Strigidae, Lybiidae, Ramphastidae, Capitonidae, and Phasianidae (Figure 3). Overall, avian susceptibility to malaria exhibited a moderate degree of phylogenetic conservatism (Pagel’s *λ* = 0.47, *P* ¡ 0.001).

**Fig 3.**
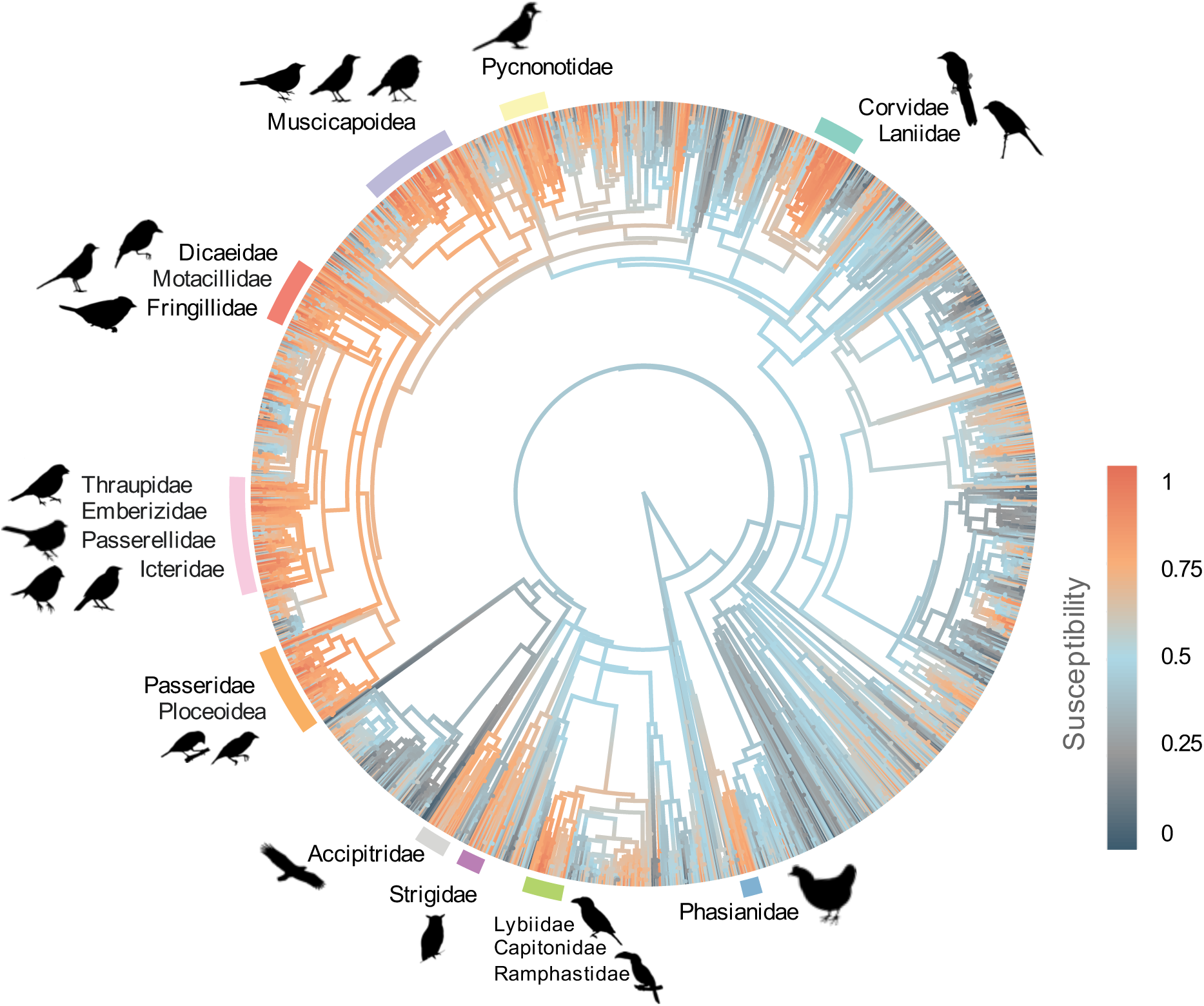
Susceptibility prediction of avian malaria in birds using interpretable machine learning models and its mapping onto the avian phylogenetic tree. A gradient from gray to orange represents increasing host susceptibility, with families exhibiting high susceptibility to avian malaria highlighted with colored blocks in the outer ring.

Notably, 13 threatened species showed relatively high susceptibility (*≥* 50%), including one critically endangered species: the Blue-banded Kingfisher (*Alcedo euryzona*); four endangered species: the Grey-cheeked Bulbul (*Alophoixus bres*), Javan Leafbird (*Chloropsis cochinchinensis*), Yellow Cardinal (*Gubernatrix cristata*), and Fernwren (*Oreoscopus gutturalis*); as well as eight vulnerable species (Table 2).

**Table 2.** Endangered bird species predicted to be susceptible to avian malaria by the machine learning model. This table presents bird species predicted to be highly susceptible (probability >0.5) and classified as vulnerable or at <u>higher risk of extinction.</u>

| Common name | Scientific name | Order | Family | Threat status | Susceptibility |
| --- | --- | --- | --- | --- | --- |
| Blue-banded Kingfisher | *Alcedo euryzona* | Coraciiformes | Alcedinidae | Critically Endangered | 0.51 |
| Grey-cheeked Bulbul | *Alophoixus bres* | Passeriformes | Pycnonotidae | Endangered | 0.85 |
| Javan Leafbird | *Chloropsis cochinchinensis* | Passeriformes | Chloropseidae | Endangered | 0.68 |
| Yellow Cardinal | *Gubernatrix cristata* | Passeriformes | Thraupidae | Endangered | 0.62 |
| Fernwren | *Oreoscopus gutturalis* | Passeriformes | Acanthizidae | Endangered | 0.50 |
| Southern Grey Shrike | *Lanius meridionalis* | Passeriformes | Laniidae | Vulnerable | 0.82 |
| Pinyon Jay | *Gymnorhinus cyanocephalus* | Passeriformes | Corvidae | Vulnerable | 0.81 |
| Opal-crowned Manakin | *Lepidothrix iris* | Passeriformes | Pipridae | Vulnerable | 0.70 |
| Red-throated Piping Guan | *Pipile cujubi* | Galliformes | Cracidae | Vulnerable | 0.69 |
| White-bellied Parrot | *Pionites leucogaster* | Psittaciformes | Psittacidae | Vulnerable | 0.66 |
| Crimson-bellied Parakeet | *Pyrrhura lepida* | Psittaciformes | Psittacidae | Vulnerable | 0.62 |
| Rondonia Warbling Antbird | *Hypocnemis ochrogyna* | Passeriformes | Thamnophilidae | Vulnerable | 0.60 |
| Green-winged Trumpeter | *Psophia viridis* | Gruiformes | Psophiidae | Vulnerable | 0.58 |

### 3.3 Drivers of global infection probability

According to the F1 score, the Xgboost model exhibited the best fit to the observed data and the strongest predictive capacity according to the mean f1 score (Table S3). Hence, we report results from this model unless stated otherwise. The best model distinguished highly susceptible bird species from less susceptible ones with fair accuracy (F1 score = 0.72 *±* 0.0003 *SE*, recall = 0.83 *±* 0.0004 *SE*, AUC = 0.70 *±* 0.0007 *SE*). According to the mean Shapley values, some top features for describing avian malaria infection probability were foraging (i.e. fruit in diet, ground foraging state, and invertebrates in diet), life history (i.e. tarsus length, beak length, and bodymass), behavior (i.e. HWI and migration distance), evolution (i.e. diversification rate and phylogenetic category), biogeography (i.e. range size, habitat breath), and human tolerance index (Figure 4). Diversification rate and range size were the first two most important predictors of susceptibility to avian malaria (SHAP value 0.5 and 0.47, separately; Figure 4). Generally, bird species with larger ranges, reduced flight or non-migratory behavior, longer ground-foraging times, greater body mass, intermediate beak length, and moderate proportion of invertebrates in diet were more susceptible to malaria infections (Figure 4). Conversely, species with lower proportion of fruit in diet or moderate habitat breadth were generally less susceptible.(Figure 4). From an evolutionary viewpoint, hosts with a moderate diversification rate tend to have the highest level of malaria infection.

**Fig 4.**
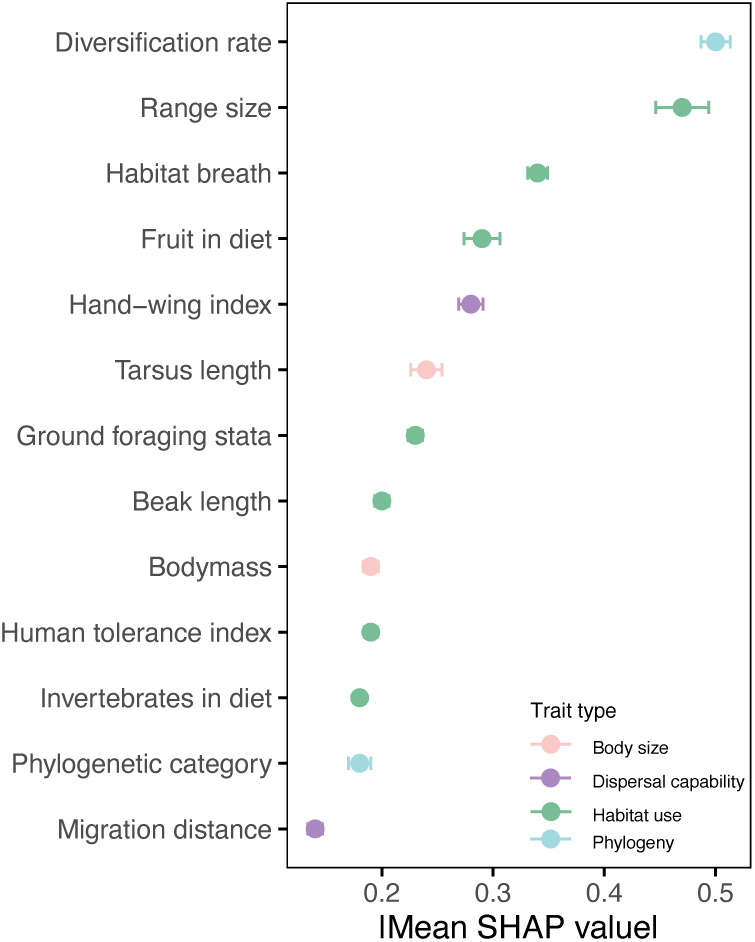
Key features associated with avian malaria susceptibility predicted by interpretable machine learning models. Only features with an absolute SHAP value greater than 0.1 are shown, ranked in descending order of feature importance.

## 4 DISCUSSION

In this study, we integrated the largest dataset to date of species-level traits and avian malaria parasite infection status, encompassing malaria infection data for 72,406 individual birds across 2,539 species. To enhance our understanding of the pattern of bird susceptibility to avian malaria at global scales, we developed a methodological framework of IML algorithms to detect host species susceptible to avian malaria among under-sampled species, revealing how hosts’ ecological and evolutionary traits jointly drive interspecific differences in susceptibility. These findings provide a novel perspective on the complex mechanisms of avian malaria transmission and offer optimized strategies for effectively monitoring and controlling the risk of Plasmodium spread in birds.

### 4.1 Ecological drivers of species-level avian malaria infections

We provide a modeling framework for testing a set of hypotheses and relevant factors for specieslevel avian malaria infections and showed their trends.

In terms of diffusion capacity, we found birds that have migratory behaviors as well as better flight capabilities were less susceptible to avian malaria (Figure 5). These results are consistent with the theoretical expectations of the “migratory escape hypothesis” [56], which posits that migratory birds reduce infection rates by avoiding infected individuals or contaminated habitats during migration. They also align with the predictions of the ”migratory culling hypothesis” [10], which suggests that migration consumes substantial energy, causing infected individuals to fail in completing migration and thus be removed from the population, leading to lower infection levels in migratory groups. However, this finding contradicts the ”migratory exposure hypothesis” [3, 45], which argues that migratory birds encounter a diversity of habitats, increasing pathogen exposure and thereby accumulating higher infection rates. Birds with wide geographic distributions exhibit higher susceptibility to avian malaria (Fig. 5), possibly because they have increased opportunities to come into contact with malaria parasites from different geographic populations.

**Fig 5.**
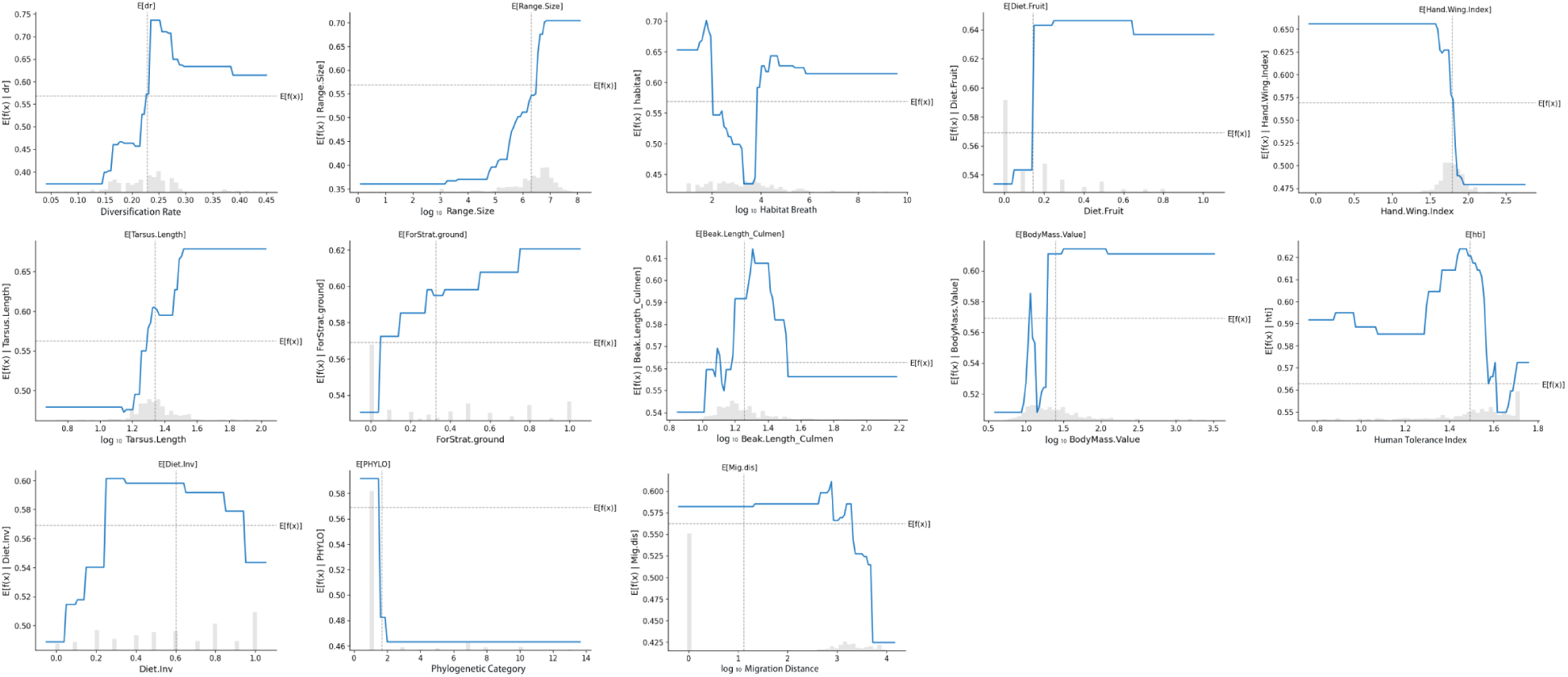
Partial dependence plots showing the effect of each feature on host susceptibility to avian malaria parasites. Blue lines represent the average effect across model iterations. Histograms display the distribution of predictor variables of the training set. Only features with an absolute SHAP value greater than 0.1 % are shown.

In habitat utilization, birds spending longer time foraging on the ground and understory strata are likely to be susceptible rather than those foraging at the canopy (Figure 5). The abundance of vector insects varies in different vertical strata, thereby affecting the exposure risk of hosts that prefer different forest layers [11, 29, 59]. Invertebrate vector hosts of *Plasmodium* (*Culex* spp. and *Aedes spp.*) are known to feed primarily at the ground level, thus increasing the prevalence of species foraging at ground strata [11, 29, 63]. Diet preferences may indirectly influence susceptibility by affecting the exposure to vectors during feeding. We found that susceptibility reached the highest when the proportions of fruits, invertebrates, and seeds in the diet were each at a moderate level (Figure 4). These may indicate that omnivorous birds are more susceptible to malaria parasites than those that feed exclusively on one type of food. In addition, birds with moderate habitat breadths tend to have a lower probability of contracting malaria parasites, whereas low or high habitat breadths correspond to higher probabilities of infection. This may reflect a trade-off of escape and exposure. Hosts with narrow habitat breadths may undergo long-term co-evolution with their malaria parasites without succumbing to death, thus exhibiting high susceptibility and suitability. If hosts leave an infected habitat and move to a new one, their prevalence may decline according to the ”migratory escape hypothesis”. However, the expansion of habitat types also increases the risk of encountering novel pathogens. Different populations of the same bird species inhabiting distinct environments may therefore experience varying levels of exposure risk, ultimately contributing to the accumulation of infections at the species level[28, 47].

It is generally recognized that human activities exacerbate the transmission of parasites[28, 47]. However, we found birds that can better tolerate human activity are less susceptible (Figure 5). A worldwide meta-analysis revealed that in areas influenced by human activity, the abundance and diversity of mosquitoes are generally lower compared to natural habitats [68], which may reduce the chances of hosts encountering malaria parasites. Besides, birds with a low HTI tend to have a decreasing population trend and be threatened [60]. High susceptibility to malaria may partly account for this phenomenon, underscoring the importance of targeted attention and conservation of these bird species.

For life-history traits, we found larger bird species tend to have a higher probability of infection (Figure 5). This pattern contradicts the expectations of ”life history theory”, which predicts that species with fast life history strategies (short lifespan, large clutch size, and early sexual maturity) are more susceptible [90]. This could be due to several reasons. First, larger body mass usually corresponds to a larger amount of bare skin exposed, making them more conspicuous and exposed in the environment, and thus more susceptible to mosquito attacks. Second, bird species with a larger body mass often emit more carbon dioxide, increasing the biting rate of mosquitoes [22, 36]. Third, species with larger body sizes have more developed immune systems. Avian heterophils scale steeply with their body mass[77]. Heterophils are phagocytes that protect predominantly against microparasites. Thus, larger species may be less likely to die from malaria and present a high prevalence.

It should be noted that the prevalence of species fluctuates, especially when the sample amount is small. To ensure the reliability of our models, species with sample sizes greater than 30 were defined as well-sampled species, and classification models were used to reduce fluctuations in prevalence, thereby improving analytical robustness. Notably, in our dataset, 1,574 bird species had fewer than 10 sampled individuals (61.9% of all species), indicating that a large number of bird species worldwide remain under-sampled, with their true infection status still uncertain. We anticipate that with increased future sampling efforts of avian malaria, infection risk assessments will become more accurate. In addition, ecological drivers may vary across temporal and spatial scales [48, 95]. Our study aimed to explore species-level infection risk drivers, thus employing species-level biogeographical factors such as range size and habitat breadth. For instance, we could collect morphometric data at the individual level by banding birds, allowing us to consider the impact of intra-specific trait variability (ITV) on functional diversity, and thus help elucidate the true dynamics and functionality of communities [14, 98]. We could also use drones to obtain high-resolution geographic information. Besides, more effort should be invested in integrating micro-level variables (e.g. genome features and mutation under laboratory conditions) and macro-level variables driving pathogen transmission, to more comprehensively assess emerging infectious diseases.

### 4.2 Relationship between evolutionary history and susceptibility to pathogen

Variation in the prevalence of haemosporidian parasites across host clades has been linked to the phylogenetic position of the hosts [6, 16]. Phylogenetic reconstruction revealed that avian malaria prevalence is highly variable within and among host clades. Susceptibility may has independently evolved multiple times and was prevalent within the order Passeriformes (Figure 3). We observed a medium phylogenetic signal in predicted probabilities, suggesting susceptibility to be phylogenetically conserved. Meanwhile, IML model showed that the host diversification rate plays an important role in malaria prevalence (Figure 4). These findings support the hypothesis that host evolutionary processes affect host susceptibility to pathogens and reject the ‘lability hypothesis’. This may because phylogenetically closed bird species tend to have conserved immune systems thus exhibiting similar prevalence [65].

Birds with moderate diversification rates are more susceptible to avian malaria, which contrasts with a previous computational model study [21]. Host species may respond to selective pressure from parasites through rapid diversification. Species with a rapid diversification rate are typically apical in phylogenies and considered to have higher ecological niche lability [103]. There may not be enough time for malaria parasites to break through their immune defenses, thus appearing to have a lower susceptibility. For those having low diversification rates, which are known as ’specialists’, they may have experienced long-term co-evolution with their vectors and pathogens, maintaining a stable state with relatively low infection rates. These results indicate that parasite-mediated selection may drive host diversification. We are unsure whether this is widely applicable and look forward to testing this hypothesis in other taxa.

### 4.3 Application of machine learning framework in ecology

Our model provides a list of highly susceptible hosts that some are consistent with previous studies(e.g., Parulidae, Turdidae, and Conopophagidae) [24, 75], while others are revealed for the first time in the global scale (e.g., Strigidae, Accipitridae, and Muscicapidea). predicted high-susceptibility species currently show low infection prevalence, such as the magpie tanager (*Cissopis leverianus*) and the common linnet (*Carduelis cannabina*). These findings can serve as a guide for potential vital hosts and reservoirs of avian malaria for future sampling.

Despite DL models for tabular data problems currently receiving increasing attention, the ML algorithms based on gradient-boosted decision trees (GBDT) seem to be a stronger solution for our dataset. This might be due to several reasons: neural networks may prefer overly smooth solutions, uninformative features might impact Multi-Layer Perceptrons (MLP)-like neural networks more significantly, or the data might not be invariant to rotation, affecting the learning process [34]. We look forward to the development of end-to-end deep learning algorithms that are advantageous for solving tabular problems in the future.

The efficiency, predictive power, and ability to model complex nonlinear patterns with minimal assumptions make ML/DL algorithms a suitable option for our global-scale large dataset of avian malaria parasites [27, 75]. Moreover, the progress in data science over the past decade means that there is no need to consider these algorithms as ’black boxes’ [27]. IML garners novel insights into the model structure that can help ecologists discover possible relationships behind the data and propose new hypotheses to be tested.

## Data Accessibility Statement

Bird infection data used in this study are freely available via MalAvi (http://130.235.244.92/Malavi/) and in Table S1 of Fecchio et al. (2021) [24]. The list of bird species predicted to have high susceptibility to avian malaria (probability *≥* 0.7) is provided in Table S4.

