## supplemental File for "What kind of birds are more susceptible to avian malaria? A global analysis based on interpretable machine learning approach"

**Xinyi Wang**[a,b,c], **Swapnil Mishra**[c,*], **Lu Dong**[a,*]
[a]MOE Key Laboratory for Biodiversity Science and Ecological Engineering, College of Life Sciences, Beijing Normal University, Beijing, China
[b]National Health Commission Key Laboratory of Parasitic Disease Control and Prevention, Jiangsu Provincial Key Laboratory on Parasite and Vector Control Technology, Jiangsu Institute of Parasitic Diseases, Wuxi, China
[c]Saw Swee Hock School of Public Health, National University of Singapore, Singapore, Singapore

### Supplementary Methods

We set the minimum sample size ($n_{min}$) and quantile threshold (q) for susceptibility classification and performed sensitivity tests to evaluate their influence on model accuracy and robustness. Bird species were divided into well-sampled and under-sampled groups. Host traits of well-sampled species were used as model inputs, while the susceptibility of under-sampled species was predicted using classification models. $n_{min}$ denotes the minimum number of individuals required for a species to be considered well-sampled. A larger $n_{min}$ increases the reliability of prevalence estimates but reduces the number of species available for training, potentially lowering model accuracy. The quantile threshold (q) determined the cutoffs for susceptibility classes. For the binary classification model, species in the top q quantile of prevalence were assigned to the high-susceptibility class (class 1), those in the bottom q quantile to the low-susceptibility class (class 0), while species in the middle range were excluded. For the ternary classification model, the middle range was instead defined as the medium-susceptibility class.

We first compared binary and ternary models. The binary model outperformed the ternary model in both F1 score and precision, likely because prevalence estimates are subject to sampling noise, particularly with small sample sizes. Medium-susceptibility species may shift categories if prevalence changes with larger sample sizes, reducing the reliability of the ternary model.

For the binary model, we further tested different combinations of $n_{min}$ and q. To ensure clear separation between susceptibility classes, we required that at $n = n_{min}$, the lower confidence bound of the minimum prevalence in class 1 ($p_{1min}$) exceeded the upper confidence bound of the maximum prevalence in class 0 ($p_{0max}$). Three parameter combinations satisfied this condition (Table S1). Among them, $n = n_{min}$ = 30 and q=33% yielded the largest training dataset (457 species, 58,251 individuals, representing 80.4% of all records), and this combination was selected as the final classification strategy. Two additional combinations with smaller datasets were also tested for robustness, and all three models produced similar trait importance and patterns, supporting the stability of our framework.

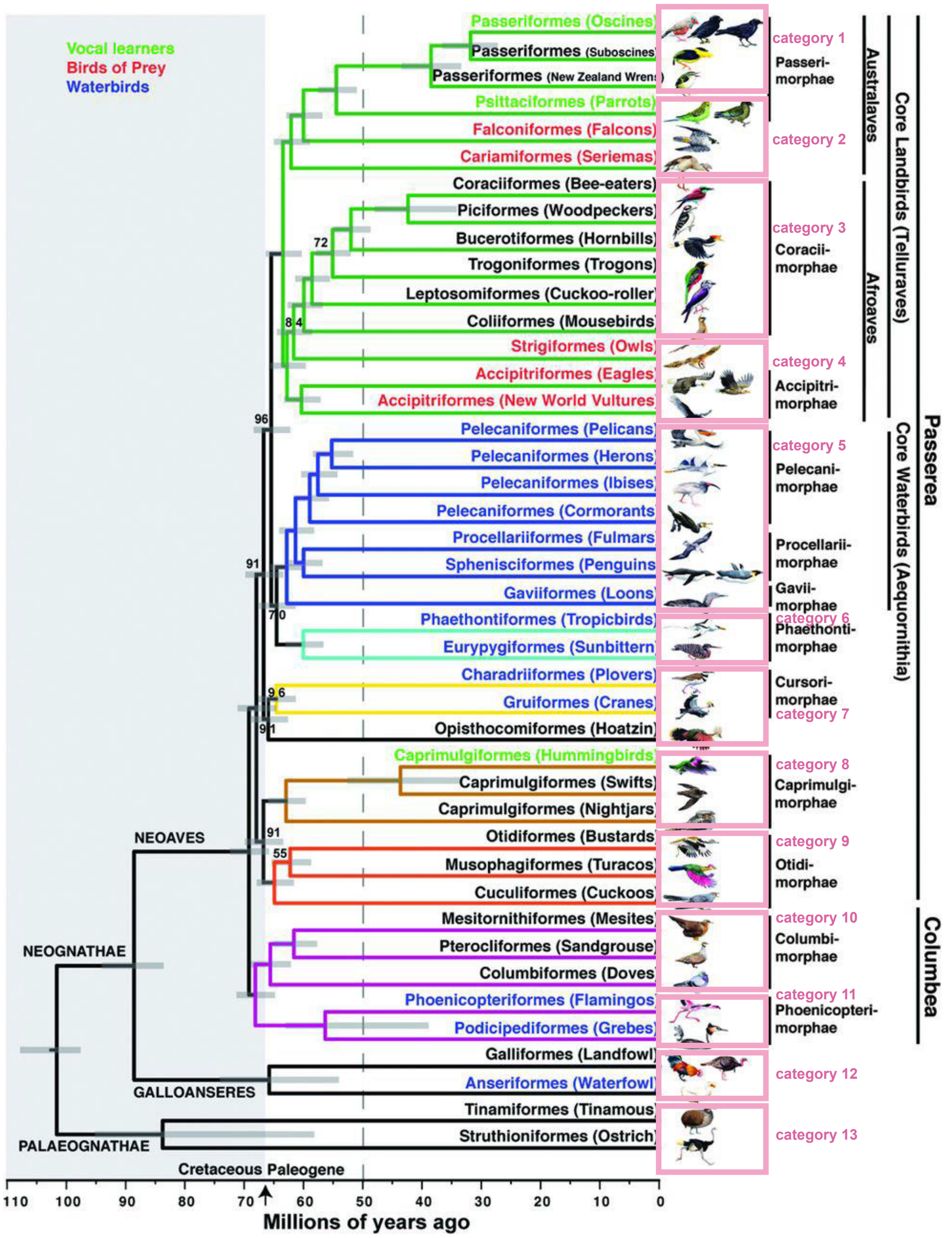


Fig S1: The 13 major avian clades defined based on a whole-genome phylogenetic tree.

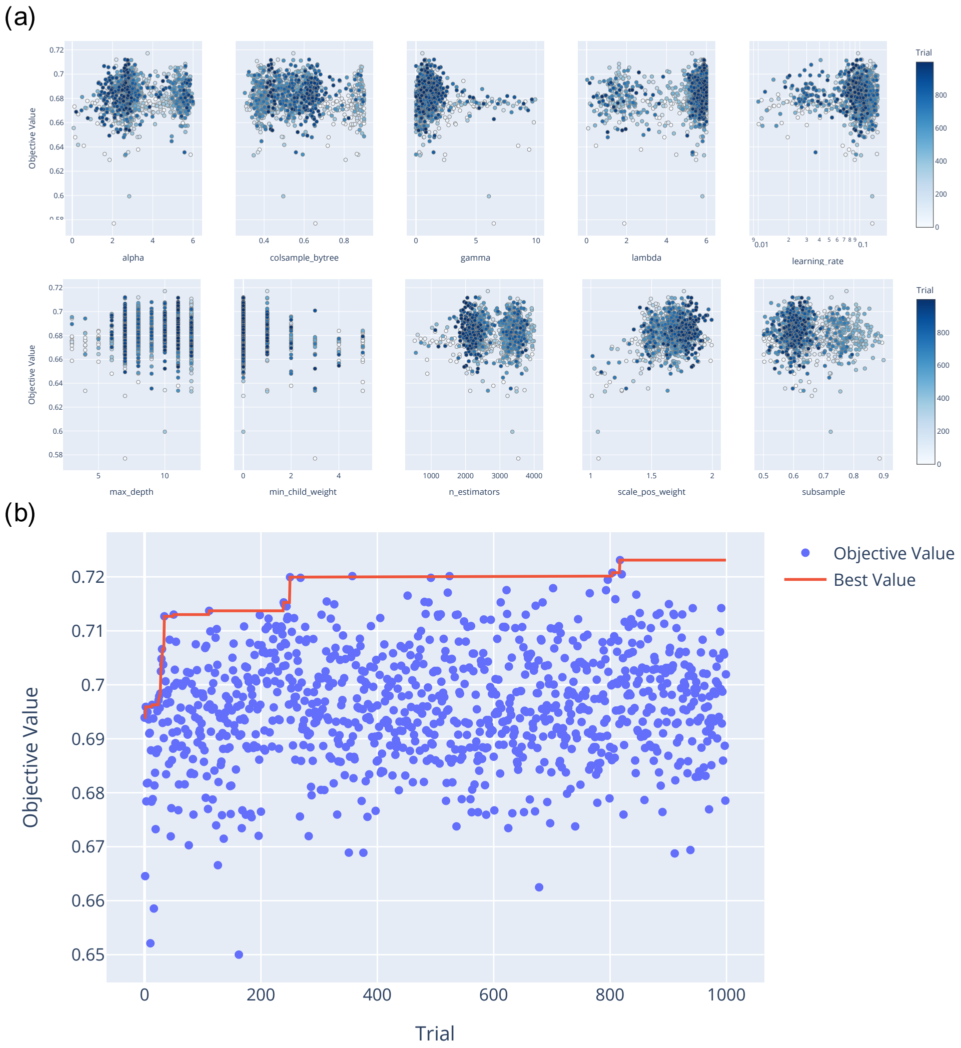


Fig S2: The results of Xgboost hyper-parameter optimization analysis using Optuna. (a) shows the distribution of hyperparameters optimized by Optuna, where each point represents a single trial, with color intensity indicating the progression of trials over time. (b) illustrates the changes in F1 score during the optimization process, with each point representing the F1 score of an individual trial. The red line indicates the highest F1 score achieved so far.

**Table S1:** Comparison of prevalence confidence intervals under different combinations of cutoff quantiles (q) and minimum sample sizes ($n_{min}$) in IML models. This table presents a comparison of prevalence confidence intervals under different combinations of minimum sample sizes and quantiles.

| Minimum sample size ($n_{min}$) | Quantile for susceptibility classification (q) | Training data size | Minimum prevalence of Class 1 ($p_{1min}$) | 95% CI lower bound of $p_{1min}$ when $n = n_{min}$ | Maximum prevalence of Class 0 ($p_{0max}$) | CI upper bound of $p_{0max}$ when $n = n_{min}$ | Meets requirement? |
|---|---|---|---|---|---|---|---|
| | 25% | 304 | 0.16 | 0.15 | 0 | 0.16 | × |
| 20 | 33% | 402 | 0.11 | 0.08 | 0 | 0.16 | × |
| | 40% | 488 | 0.08 | 0.05 | 0.02 | 0.24 | × |
| | 25% | 228 | 0.19 | 0.10 | 0.03 | 0.16 | ✓ |
| 30 | 33% | 304 | 0.16 | 0.07 | 0.05 | 0.16 | ✓ |
| | 40% | 365 | 0.13 | 0.05 | 0.07 | 0.21 | × |
| | 25% | 173 | 0.19 | 0.10 | 0.03 | 0.13 | ✓ |
| 40 | 33% | 231 | 0.12 | 0.05 | 0.05 | 0.17 | × |
| | 40% | 276 | 0.09 | 0.04 | 0.07 | 0.17 | × |

**Table S2:** Optimal hyperparameters of the XGBoost model.

| Hyperparameter | Optimal value |
|---|---|
| max_depth | 7 |
| lambda | 5.751 |
| alpha | 2.350 |
| min_child_weight | 0 |
| gamma | 1.304 |
| learning_rate | 0.097 |
| colsample_bytree | 0.590 |
| subsample | 0.724 |
| n_estimators | 2199 |
| scale_pos_weight | 1.790 |

**Table S3:** Average performance evaluation of XGBoost, TabR, and TabDDPM models in predicting avian malaria susceptibility.

| Name | Mean F1 score | Mean Recall | Mean AUC |
|---|---|---|---|
| XGBoost | 0.72 | 0.83 | 0.70 |
| TabR | 0.61 | 0.63 | 0.66 |
| TabDDPM | 0.57 | 0.55 | 0.58 |

**Table S4:** Bird species with high susceptibility to malaria predicted by interpretable machine learning models

| Species | Susceptibility | Species | Susceptibility |
|---|---|---|---|
| *Cissopis leverianus* | 0.98 | *Dendroica striata* | 0.82 |
| *Fringilla montifringilla* | 0.98 | *Empidonax alnorum* | 0.82 |
| *Icterus spurius* | 0.98 | *Euphonia hirundinacea* | 0.82 |
| *Saltator atricollis* | 0.98 | *Fluvicola albiventer* | 0.82 |
| *Zoothera citrina* | 0.98 | *Hylophilus pectoralis* | 0.82 |
| *Carduelis cannabina* | 0.97 | *Indicator exilis* | 0.82 |
| *Creatophora cinerea* | 0.97 | *Iole olivacea* | 0.82 |
| *Erythropygia galactotes* | 0.97 | *Lanius meridionalis* | 0.82 |
| *Gnorimopsar chopi* | 0.97 | *Lipaugus vociferans* | 0.82 |
| *Myrmecocichla cinnamomeiventris* | 0.97 | *Megalaima henricii* | 0.82 |
| *Ploceus nigricollis* | 0.97 | *Megascops watsonii* | 0.82 |
| *Thescelocichla leucopleura* | 0.97 | *Monticola gularis* | 0.82 |
| *Turdus pelios* | 0.97 | *Muscicapa sibirica* | 0.82 |
| *Turdus pilaris* | 0.97 | *Pachyramphus minor* | 0.82 |
| *Cyanerpes caeruleus* | 0.96 | *Passer diffusus* | 0.82 |
| *Cypsnagra hirundinacea* | 0.96 | *Poospiza nigrorufa* | 0.82 |
| *Lamprotornis chalybaeus* | 0.96 | *Prionochilus maculatus* | 0.82 |
| *Lybius vieilloti* | 0.96 | *Quelea quelea* | 0.82 |
| *Pipilo maculatus* | 0.96 | *Serinus sulphuratus* | 0.82 |
| *Ploceus cucullatus* | 0.96 | *Sturnus unicolor* | 0.82 |
| *Ploceus intermedius* | 0.96 | *Tricholaema diademata* | 0.82 |
| *Pycnonotus jocosus* | 0.96 | *Turdus serranus* | 0.82 |
| *Sialia sialis* | 0.96 | *Aegolius acadicus* | 0.81 |
| *Sturnella militaris* | 0.96 | *Bucco tamatia* | 0.81 |
| *Thryothorus coraya* | 0.96 | *Corythopis torquatus* | 0.81 |
| *Turdus hauxwelli* | 0.96 | *Cyanopica cyanus* | 0.81 |
| *Carduelis psaltria* | 0.95 | *Dendrocolaptes picumnus* | 0.81 |
| *Carpodacus erythrinus* | 0.95 | *Dendroica pinus* | 0.81 |
| *Cinclus cinclus* | 0.95 | *Emberiza pusilla* | 0.81 |
| *Cyphorhinus arada* | 0.95 | *Emberiza rutila* | 0.81 |
| *Emberizoides herbicola* | 0.95 | *Estrilda caerulescens* | 0.81 |
| *Euplectes hordeaceus* | 0.95 | *Euphonia chlorotica* | 0.81 |
| *Lanius excubitor* | 0.95 | *Gallus gallus* | 0.81 |
| *Nectarinia cuprea* | 0.95 | *Gymnorhinus cyanocephalus* | 0.81 |
| *Nigrita bicolor* | 0.95 | *Lagonosticta rara* | 0.81 |
| *Oenanthe pleschanka* | 0.95 | *Manorina melanocephala* | 0.81 |
| *Passerella iliaca* | 0.95 | *Megascops kennicottii* | 0.81 |

(Continued) Bird species with high susceptibility to malaria predicted by interpretable machine learning models

| Species | Susceptibility | Species | Susceptibility |
|---|---|---|---|
| *Petronia petronia* | 0.95 | *Parula pitiayumi* | 0.81 |
| *Pycnonotus goiavier* | 0.95 | *Phainopepla nitens* | 0.81 |
| *Spermophaga haematina* | 0.95 | *Phoenicurus hodgsoni* | 0.81 |
| *Taraba major* | 0.95 | *Pitangus sulphuratus* | 0.81 |
| *Turdoides plebejus* | 0.95 | *Psarocolius angustifrons* | 0.81 |
| *Turdus fumigatus* | 0.95 | *Ramphocelus passerinii* | 0.81 |
| *Turdus ignobilis* | 0.95 | *Saltator coerulescens* | 0.81 |
| *Zoothera sibirica* | 0.95 | *Saltator maxillosus* | 0.81 |
| *Anthreptes collaris* | 0.94 | *Sclerurus mexicanus* | 0.81 |
| *Attila spadiceus* | 0.94 | *Spermophaga ruficapilla* | 0.81 |
| *Cossypha niveicapilla* | 0.94 | *Tachyphonus luctuosus* | 0.81 |
| *Cyanocitta stelleri* | 0.94 | *Tangara larvata* | 0.81 |
| *Loxia leucoptera* | 0.94 | *Tangara punctata* | 0.81 |
| *Nectarinia chloropygia* | 0.94 | *Thlypopsis sordida* | 0.81 |
| *Passerina caerulea* | 0.94 | *Trachyphonus darnaudii* | 0.81 |
| *Pheucticus ludovicianus* | 0.94 | *Tricholaema hirsuta* | 0.81 |
| *Pheucticus melanocephalus* | 0.94 | *Tricholaema lacrymosa* | 0.81 |
| *Pooecetes gramineus* | 0.94 | *Uraeginthus angolensis* | 0.81 |
| *Psarocolius decumanus* | 0.94 | *Vidua macroura* | 0.81 |
| *Saxicola caprata* | 0.94 | *Cephalopterus ornatus* | 0.8 |
| *Turdus assimilis* | 0.94 | *Chlorestes notata* | 0.8 |
| *Turdus libonyanus* | 0.94 | *Circus cyaneus* | 0.8 |
| *Turdus subalaris* | 0.94 | *Crotophaga major* | 0.8 |
| *Turdus viscivorus* | 0.94 | *Cyanocompsa parellina* | 0.8 |
| *Agelaioides badius* | 0.93 | *Dendragapus canadensis* | 0.8 |
| *Ammodramus savannarum* | 0.93 | *Dendrocopos syriacus* | 0.8 |
| *Andropadus gracilirostris* | 0.93 | *Dubusia taeniata* | 0.8 |
| *Andropadus importunus* | 0.93 | *Entomyzon cyanotis* | 0.8 |
| *Anthus rufulus* | 0.93 | *Eremophila alpestris* | 0.8 |
| *Anthus similis* | 0.93 | *Lagonosticta rufopicta* | 0.8 |
| *Chlorocichla flaviventris* | 0.93 | *Lagonosticta senegala* | 0.8 |
| *Corvus brachyrhynchos* | 0.93 | *Lanius tephronotus* | 0.8 |
| *Cyanocorax cyanomelas* | 0.93 | *Mandingoa nitidula* | 0.8 |
| *Cyanocorax morio* | 0.93 | *Melocichla mentalis* | 0.8 |
| *Eurocephalus rueppelli* | 0.93 | *Mirafra africana* | 0.8 |
| *Hypogramma hypogrammicum* | 0.93 | *Myrmoborus leucophrys* | 0.8 |
| *Lamprospiza melanoleuca* | 0.93 | *Neocichla gutturalis* | 0.8 |
| *Lanius ludovicianus* | 0.93 | *Ortygospiza atricollis* | 0.8 |

(Continued) Bird species with high susceptibility to malaria predicted by interpretable machine learning models

| Species | Susceptibility | Species | Susceptibility |
|---|---|---|---|
| *Nectarinia verticalis* | 0.93 | *Pachyramphus polychopterus* | 0.8 |
| *Oenanthe oenanthe* | 0.93 | *Ploceus rubiginosus* | 0.8 |
| *Paradoxornis webbianus* | 0.93 | *Ploceus superciliosus* | 0.8 |
| *Petronia superciliaris* | 0.93 | *Pogoniulus bilineatus* | 0.8 |
| *Phyllastrephus cerviniventris* | 0.93 | *Pygiptila stellaris* | 0.8 |
| *Phyllastrephus terrestris* | 0.93 | *Schistocichla leucostigma* | 0.8 |
| *Piranga bidentata* | 0.93 | *Sicalis flaveola* | 0.8 |
| *Saxicola torquatus* | 0.93 | *Sicalis olivascens* | 0.8 |
| *Schistochlamys melanopis* | 0.93 | *Sporophila nigricollis* | 0.8 |
| *Sericornis frontalis* | 0.93 | *Terpsiphone viridis* | 0.8 |
| *Spizella arborea* | 0.93 | *Thamnophilus torquatus* | 0.8 |
| *Turdoides sharpei* | 0.93 | *Trogon curucui* | 0.8 |
| *Amblyospiza albifrons* | 0.92 | *Tyranneutes stolzmanni* | 0.8 |
| *Anaplectes rubriceps* | 0.92 | *Veniliornis spilogaster* | 0.8 |
| *Arremon brunneinucha* | 0.92 | *Zosterops erythropleurus* | 0.8 |
| *Bradornis pallidus* | 0.92 | *Chlorostilbon mellisugus* | 0.79 |
| *Carduelis spinus* | 0.92 | *Cisticola brachypterus* | 0.79 |
| *Catharus aurantiirostris* | 0.92 | *Cisticola juncidis* | 0.79 |
| *Catharus occidentalis* | 0.92 | *Dryoscopus cubla* | 0.79 |
| *Chondestes grammacus* | 0.92 | *Emberiza cioides* | 0.79 |
| *Furnarius rufus* | 0.92 | *Estrilda melanotis* | 0.79 |
| *Habia fuscicauda* | 0.92 | *Estrilda perreini* | 0.79 |
| *Icterus parisorum* | 0.92 | *Formicivora rufa* | 0.79 |
| *Luscinia calliope* | 0.92 | *Forpus xanthopterygius* | 0.79 |
| *Luscinia svecica* | 0.92 | *Hylophilus poicilotis* | 0.79 |
| *Lybius leucocephalus* | 0.92 | *Indicator indicator* | 0.79 |
| *Macronyx croceus* | 0.92 | *Lagopus lagopus* | 0.79 |
| *Miliaria calandra* | 0.92 | *Lamprotornis chloropterus* | 0.79 |
| *Oenanthe isabellina* | 0.92 | *Lanius senator* | 0.79 |
| *Ploceus velatus* | 0.92 | *Limnothlypis swainsonii* | 0.79 |
| *Saltator grossus* | 0.92 | *Melanocharis nigra* | 0.79 |
| *Tersina viridis* | 0.92 | *Muscicapa dauurica* | 0.79 |
| *Thraupis palmarum* | 0.92 | *Myrmeciza melanoceps* | 0.79 |
| *Turdus naumanni* | 0.92 | *Oporornis philadelphia* | 0.79 |
| *Turdus rubrocanus* | 0.92 | *Passerina amoena* | 0.79 |
| *Urocissa erythrorhyncha* | 0.92 | *Phyllastrephus xavieri* | 0.79 |
| *Anthus leucophrys* | 0.91 | *Pulsatrix perspicillata* | 0.79 |
| *Anthus spinoletta* | 0.91 | *Ramphocaenus melanurus* | 0.79 |

(Continued) Bird species with high susceptibility to malaria predicted by interpretable machine learning models

| Species | Susceptibility | Species | Susceptibility |
|---|---|---|---|
| *Aphelocoma ultramarina* | 0.91 | *Rhytipterna simplex* | 0.79 |
| *Arremon flavirostris* | 0.91 | *Sporophila torqueola* | 0.79 |
| *Arremonops conirostris* | 0.91 | *Sporopipes frontalis* | 0.79 |
| *Arremonops rufivirgatus* | 0.91 | *Stephanophorus diadematus* | 0.79 |
| *Cacicus cela* | 0.91 | *Sylvia hortensis* | 0.79 |
| *Charitospiza eucosma* | 0.91 | *Tachyphonus rufiventer* | 0.79 |
| *Cisticola chiniana* | 0.91 | *Tangara velia* | 0.79 |
| *Cisticola natalensis* | 0.91 | *Turdoides reinwardii* | 0.79 |
| *Copsychus saularis* | 0.91 | *Turdus poliocephalus* | 0.79 |
| *Cyanerpes cyaneus* | 0.91 | *Veniliornis affinis* | 0.79 |
| *Cyanocorax cyanopogon* | 0.91 | *Vireo plumbeus* | 0.79 |
| *Hippolais pallida* | 0.91 | *Agriornis lividus* | 0.78 |
| *Lanius cristatus* | 0.91 | *Donacospiza albifrons* | 0.78 |
| *Mimus patagonicus* | 0.91 | *Dryocopus martius* | 0.78 |
| *Molothrus bonariensis* | 0.91 | *Empidonax flaviventris* | 0.78 |
| *Nucifraga caryocatactes* | 0.91 | *Eremopterix leucotis* | 0.78 |
| *Petronia pyrgita* | 0.91 | *Estrilda troglodytes* | 0.78 |
| *Pheucticus aureoventris* | 0.91 | *Ficedula narcissina* | 0.78 |
| *Pheucticus chrysogaster* | 0.91 | *Henicorhina leucosticta* | 0.78 |
| *Piranga ludoviciana* | 0.91 | *Lagopus muta* | 0.78 |
| *Ploceus ocularis* | 0.91 | *Microbates collaris* | 0.78 |
| *Ploceus xanthops* | 0.91 | *Monasa nigrifrons* | 0.78 |
| *Pyrrhocorax pyrrhocorax* | 0.91 | *Oporornis tolmiei* | 0.78 |
| *Sylvia nisoria* | 0.91 | *Phoenicurus moussieri* | 0.78 |
| *Vermivora celata* | 0.91 | *Phyllastrephus flavostriatus* | 0.78 |
| *Vireo flavifrons* | 0.91 | *Phylloscopus fuscatus* | 0.78 |
| *Ammodramus aurifrons* | 0.9 | *Ploceus luteolus* | 0.78 |
| *Arremon aurantiirostris* | 0.9 | *Pogoniulus leucomystax* | 0.78 |
| *Attila cinnamomeus* | 0.9 | *Querula purpurata* | 0.78 |
| *Cercomela familiaris* | 0.9 | *Sclerurus rufigularis* | 0.78 |
| *Cisticola galactotes* | 0.9 | *Sicalis citrina* | 0.78 |
| *Cyanocorax yncas* | 0.9 | *Sicalis uropygialis* | 0.78 |
| *Dicaeum cruentatum* | 0.9 | *Sylvietta virens* | 0.78 |
| *Lullula arborea* | 0.9 | *Tangara xanthogastra* | 0.78 |
| *Lybius torquatus* | 0.9 | *Thamnophilus atrinucha* | 0.78 |
| *Molothrus rufoaxillaris* | 0.9 | *Thamnophilus nigrocinereus* | 0.78 |
| *Monticola rufocinereus* | 0.9 | *Wilsonia pusilla* | 0.78 |
| *Monticola saxatilis* | 0.9 | *Accipiter striatus* | 0.77 |

(Continued) Bird species with high susceptibility to malaria predicted by interpretable machine learning models

| Species | Susceptibility | Species | Susceptibility |
|---|---|---|---|
| *Myadestes townsendi* | 0.9 | *Ammodramus leconteii* | 0.77 |
| *Nectarinia aspasia* | 0.9 | *Aquila wahlbergi* | 0.77 |
| *Nectarinia cyanolaema* | 0.9 | *Arachnothera longirostra* | 0.77 |
| *Nigrita canicapillus* | 0.9 | *Batara cinerea* | 0.77 |
| *Pipraeidea melanonota* | 0.9 | *Bombycilla japonica* | 0.77 |
| *Pomatostomus temporalis* | 0.9 | *Calliphlox amethystina* | 0.77 |
| *Poospiza melanoleuca* | 0.9 | *Celeus flavescens* | 0.77 |
| *Serinus mozambicus* | 0.9 | *Cinclus leucocephalus* | 0.77 |
| *Sturnella magna* | 0.9 | *Cisticola erythrops* | 0.77 |
| *Sylvia cantillans* | 0.9 | *Crateroscelis murina* | 0.77 |
| *Turdus olivaceus* | 0.9 | *Cryptospiza reichenovii* | 0.77 |
| *Zosterops japonicus* | 0.9 | *Emberiza cabanisi* | 0.77 |
| *Aphelocoma californica* | 0.89 | *Emberiza melanocephala* | 0.77 |
| *Baeopogon indicator* | 0.89 | *Emberiza tahapisi* | 0.77 |
| *Cacicus haemorrhous* | 0.89 | *Erithacus akahige* | 0.77 |
| *Copsychus malabaricus* | 0.89 | *Eubucco richardsoni* | 0.77 |
| *Corvus corax* | 0.89 | *Hylia prasina* | 0.77 |
| *Coryphospingus cucullatus* | 0.89 | *Hylophilus amaurocephalus* | 0.77 |
| *Coturnix delegorguei* | 0.89 | *Lonchura fringilloides* | 0.77 |
| *Estrilda melpoda* | 0.89 | *Meliphaga analoga* | 0.77 |
| *Euphonia minuta* | 0.89 | *Oriolus auratus* | 0.77 |
| *Myiarchus cinerascens* | 0.89 | *Phasianus colchicus* | 0.77 |
| *Nectarinia seimundi* | 0.89 | *Picoides villosus* | 0.77 |
| *Nicator chloris* | 0.89 | *Pitangus lictor* | 0.77 |
| *Oenanthe hispanica* | 0.89 | *Pogoniulus chrysoconus* | 0.77 |
| *Pica pica* | 0.89 | *Pogoniulus subsulphureus* | 0.77 |
| *Pinicola enucleator* | 0.89 | *Prionops plumatus* | 0.77 |
| *Pycnonotus atriceps* | 0.89 | *Pycnonotus brunneus* | 0.77 |
| *Ramphocelus nigrogularis* | 0.89 | *Sclerurus caudacutus* | 0.77 |
| *Sialia mexicana* | 0.89 | *Sphyrapicus varius* | 0.77 |
| *Sitta neumayer* | 0.89 | *Synallaxis gujanensis* | 0.77 |
| *Tchagra minutus* | 0.89 | *Tockus alboterminatus* | 0.77 |
| *Tchagra senegalus* | 0.89 | *Xolmis cinereus* | 0.77 |
| *Turdus mupinensis* | 0.89 | *Amazilia versicolor* | 0.76 |
| *Accipiter gularis* | 0.88 | *Apalis thoracica* | 0.76 |
| *Andropadus curvirostris* | 0.88 | *Campylorhynchus zonatus* | 0.76 |
| *Baryphthengus martii* | 0.88 | *Catharus fuscater* | 0.76 |
| *Bombycilla cedrorum* | 0.88 | *Cercomacra nigrescens* | 0.76 |
| *Buteo magnirostris* | 0.88 | *Chamaeza campanisona* | 0.76 |
| *Cacicus chrysopterus* | 0.88 | *Estrilda rhodopyga* | 0.76 |

(Continued) Bird species with high susceptibility to malaria predicted by interpretable machine learning models

| Species | Susceptibility | Species | Susceptibility |
|---|---|---|---|
| *Cicinnurus regius* | 0.88 | *Laniocera hypopyrra* | 0.76 |
| *Corvus albus* | 0.88 | *Muscicapa adusta* | 0.76 |
| *Corvus corone* | 0.88 | *Myiopsitta monachus* | 0.76 |
| *Corvus orru* | 0.88 | *Nucifraga columbiana* | 0.76 |
| *Cossypha heuglini* | 0.88 | *Penelope superciliaris* | 0.76 |
| *Cossypha natalensis* | 0.88 | *Percnostola rufifrons* | 0.76 |
| *Dacnis cayana* | 0.88 | *Prionochilus percussus* | 0.76 |
| *Dacnis lineata* | 0.88 | *Rhipidura leucophrys* | 0.76 |
| *Dendroica palmarum* | 0.88 | *Strix aluco* | 0.76 |
| *Dendroica virens* | 0.88 | *Veniliornis mixtus* | 0.76 |
| *Dicaeum trigonostigma* | 0.88 | *Alethe poliocephala* | 0.75 |
| *Glaucidium perlatum* | 0.88 | *Anurolimnas viridis* | 0.75 |
| *Grallina cyanoleuca* | 0.88 | *Aramides cajanea* | 0.75 |
| *Hemithraupis flavicollis* | 0.88 | *Aratinga aurea* | 0.75 |
| *Laniarius aethiopicus* | 0.88 | *Attila rufus* | 0.75 |
| *Lanius collaris* | 0.88 | *Catharus frantzii* | 0.75 |
| *Mackenziaena severa* | 0.88 | *Conopophaga peruviana* | 0.75 |
| *Oriolus larvatus* | 0.88 | *Dendrocolaptes platy-rostris* | 0.75 |
| *Paroaria coronata* | 0.88 | *Dryocopus lineatus* | 0.75 |
| *Picoides scalaris* | 0.88 | *Emberiza tristrami* | 0.75 |
| *Plocepasser mahali* | 0.88 | *Fraseria cinerascens* | 0.75 |
| *Prunella collaris* | 0.88 | *Frederickena unduligera* | 0.75 |
| *Pycnonotus melanicterus* | 0.88 | *Larus novaehollandiae* | 0.75 |
| *Pytilia melba* | 0.88 | *Myrmeciza fortis* | 0.75 |
| *Sclateria naevia* | 0.88 | *Nilaus afer* | 0.75 |
| *Stachyris poliocephala* | 0.88 | *Parus hudsonicus* | 0.75 |
| *Tachyphonus cristatus* | 0.88 | *Phyllastrephus icterinus* | 0.75 |
| *Trachyphonus purpuratus* | 0.88 | *Ptiloris magnificus* | 0.75 |
| *Treron calvus* | 0.88 | *Ramphastos tucanus* | 0.75 |
| *Vermivora peregrina* | 0.88 | *Sitta canadensis* | 0.75 |
| *Anthus hodgsoni* | 0.87 | *Spermophaga poliogenys* | 0.75 |
| *Anthus lutescens* | 0.87 | *Turdoides rubiginosa* | 0.75 |
| *Anthus rubescens* | 0.87 | *Tyrannus savana* | 0.75 |
| *Bleda syndactylus* | 0.87 | *Bonasa umbellus* | 0.74 |
| *Carduelis magellanica* | 0.87 | *Buthraupis montana* | 0.74 |
| *Cisticola cantans* | 0.87 | *Celeus grammicus* | 0.74 |
| *Colluricincla harmonica* | 0.87 | *Ceyx pictus* | 0.74 |
| *Conirostrum cinereum* | 0.87 | *Cisticola ayresii* | 0.74 |
| *Dendrocopos medius* | 0.87 | *Cistothorus platensis* | 0.74 |
| *Dendropicos fuscescens* | 0.87 | *Dichrozona cincta* | 0.74 |

(Continued) Bird species with high susceptibility to malaria predicted by interpretable machine learning models

| Species | Susceptibility | Species | Susceptibility |
|---|---|---|---|
| *Estrilda quartinia* | 0.87 | *Donacobius atricapilla* | 0.74 |
| *Motacilla flava* | 0.87 | *Heliolais erythropterus* | 0.74 |
| *Oenanthe deserti* | 0.87 | *Hypocnemoides melanopogon* | 0.74 |
| *Parmoptila woodhousei* | 0.87 | *Icterus gularis* | 0.74 |
| *Parus cristatus* | 0.87 | *Lagonosticta rhodopareia* | 0.74 |
| *Tachyphonus surinamus* | 0.87 | *Micrastur gilvicollis* | 0.74 |
| *Campylorhynchus turdinus* | 0.86 | *Motacilla clara* | 0.74 |
| *Carduelis sinica* | 0.86 | *Myrmornis torquata* | 0.74 |
| *Cisticola fulvicapilla* | 0.86 | *Otus scops* | 0.74 |
| *Corvus albicollis* | 0.86 | *Philydor pyrrhodes* | 0.74 |
| *Cracticus nigrogularis* | 0.86 | *Piranga leucoptera* | 0.74 |
| *Cymbilaimus lineatus* | 0.86 | *Platysteira castanea* | 0.74 |
| *Dacnis flaviventer* | 0.86 | *Regulus calendula* | 0.74 |
| *Emberiza hortulana* | 0.86 | *Rupicola peruvianus* | 0.74 |
| *Eucometis penicillata* | 0.86 | *Serinus citrinelloides* | 0.74 |
| *Euplectes albonotatus* | 0.86 | *Smithornis capensis* | 0.74 |
| *Formicarius analis* | 0.86 | *Sphyrapicus thyroideus* | 0.74 |
| *Garrulax davidi* | 0.86 | *Tangara nigrocincta* | 0.74 |
| *Icterus pustulatus* | 0.86 | *Telophorus sulfureopectus* | 0.74 |
| *Loxia curvirostra* | 0.86 | *Thamnophilus caerulescens* | 0.74 |
| *Mimus triurus* | 0.86 | *Tyrannopsis sulphurea* | 0.74 |
| *Mirafra rufocinnamomea* | 0.86 | *Xolmis irupero* | 0.74 |
| *Myiarchus crinitus* | 0.86 | *Ampelion rubrocristatus* | 0.73 |
| *Parus atricapillus* | 0.86 | *Aratinga leucophthalma* | 0.73 |
| *Quiscalus mexicanus* | 0.86 | *Automolus rufipileatus* | 0.73 |
| *Tangara chilensis* | 0.86 | *Cercomacra tyrannina* | 0.73 |
| *Thraupis bonariensis* | 0.86 | *Cisticola woosnami* | 0.73 |
| *Thraupis cyanocephala* | 0.86 | *Coturnix coturnix* | 0.73 |
| *Thryothorus leucotis* | 0.86 | *Emberiza flaviventris* | 0.73 |
| *Xolmis pyrope* | 0.86 | *Empidonax minimus* | 0.73 |
| *Alophoixus bres* | 0.85 | *Harpagus bidentatus* | 0.73 |
| *Andropadus nigriceps* | 0.85 | *Hemithraupis ruficapilla* | 0.73 |
| *Athene noctua* | 0.85 | *Hypocnemis peruviana* | 0.73 |
| *Capito auratus* | 0.85 | *Illadopsis rufipennis* | 0.73 |
| *Certhia americana* | 0.85 | *Lepidothrix serena* | 0.73 |
| *Chlorophonia cyanea* | 0.85 | *Leucopternis albicollis* | 0.73 |
| *Conirostrum speciosum* | 0.85 | *Locustella luscinioides* | 0.73 |
| *Erythropygia leucophrys* | 0.85 | *Melanerpes aurifrons* | 0.73 |

(Continued) Bird species with high susceptibility to malaria predicted by interpretable machine learning models

| Species | Susceptibility | Species | Susceptibility |
|---|---|---|---|
| *Euphonia laniirostris* | 0.85 | *Melanerpes carolinus* | 0.73 |
| *Euphonia plumbea* | 0.85 | *Micrathene whitneyi* | 0.73 |
| *Euplectes macroura* | 0.85 | *Microbates cinereiventris* | 0.73 |
| *Hylocharis cyanus* | 0.85 | *Motacilla cinerea* | 0.73 |
| *Malaconotus blanchoti* | 0.85 | *Nystalus chacuru* | 0.73 |
| *Melanerpes cruentatus* | 0.85 | *Phaethornis ruber* | 0.73 |
| *Micromonacha lanceolata* | 0.85 | *Phylloscopus armandii* | 0.73 |
| *Molothrus aeneus* | 0.85 | *Pitohui ferrugineus* | 0.73 |
| *Myiozetetes similis* | 0.85 | *Pitohui kirhocephalus* | 0.73 |
| *Paroaria dominicana* | 0.85 | *Pycnonotus cyaniventris* | 0.73 |
| *Phyllastrephus albigularis* | 0.85 | *Pyroderus scutatus* | 0.73 |
| *Pica hudsonia* | 0.85 | *Ramphotrigon ruficauda* | 0.73 |
| *Psarocolius bifasciatus* | 0.85 | *Sporophila minuta* | 0.73 |
| *Ramphastos vitellinus* | 0.85 | *Tangara arthus* | 0.73 |
| *Ramphocelus bresilius* | 0.85 | *Thalurania glaucopis* | 0.73 |
| *Saltator atriceps* | 0.85 | *Tityra semifasciata* | 0.73 |
| *Saltator aurantiirostris* | 0.85 | *Zoothera gurneyi* | 0.73 |
| *Tchagra australis* | 0.85 | *Acanthiza chrysorrhoa* | 0.72 |
| *Toxostoma curvirostre* | 0.85 | *Acanthiza pusilla* | 0.72 |
| *Trogon viridis* | 0.85 | *Bubo virginianus* | 0.72 |
| *Turdus leucops* | 0.85 | *Bucco macrodactylus* | 0.72 |
| *Vidua paradisaea* | 0.85 | *Chlorornis riefferii* | 0.72 |
| *Vireo huttoni* | 0.85 | *Conopophaga roberti* | 0.72 |
| *Vireolanius leucotis* | 0.85 | *Contopus sordidulus* | 0.72 |
| *Zoothera dauma* | 0.85 | *Cynanthus latirostris* | 0.72 |
| *Alauda arvensis* | 0.84 | *Dendroica caerulescens* | 0.72 |
| *Amandava subflava* | 0.84 | *Diglossa brunneiventris* | 0.72 |
| *Amazilia fimbriata* | 0.84 | *Dives dives* | 0.72 |
| *Arremon torquatus* | 0.84 | *Elaenia obscura* | 0.72 |
| *Athene cunicularia* | 0.84 | *Gymnorhina tibicen* | 0.72 |
| *Bombycilla garrulus* | 0.84 | *Heliactin bilophus* | 0.72 |
| *Camaroptera chloronota* | 0.84 | *Hypargos niveoguttatus* | 0.72 |
| *Camaroptera undosa* | 0.84 | *Machaeropterus pyrocephalus* | 0.72 |
| *Conopophaga aurita* | 0.84 | *Machaeropterus regulus* | 0.72 |
| *Crypturellus soui* | 0.84 | *Melanodryas cucullata* | 0.72 |
| *Emberiza cia* | 0.84 | *Micrastur ruficollis* | 0.72 |
| *Euphonia chalybea* | 0.84 | *Myioparus griseigularis* | 0.72 |
| *Euphonia violacea* | 0.84 | *Phacellodomus ruber* | 0.72 |
| *Euplectes capensis* | 0.84 | *Sicalis luteola* | 0.72 |
| *Gallinula chloropus* | 0.84 | *Sporophila albogularis* | 0.72 |

(Continued) Bird species with high susceptibility to malaria predicted by interpretable machine learning models

| Species | Susceptibility | Species | Susceptibility |
|---|---|---|---|
| *Icterus cayanensis* | 0.84 | *Tangara mexicana* | 0.72 |
| *Ixos amaurotis* | 0.84 | *Tangara schrankii* | 0.72 |
| *Kaupifalco monogrammicus* | 0.84 | *Turdus infuscatus* | 0.72 |
| *Lochmias nematura* | 0.84 | *Accipiter soloensis* | 0.71 |
| *Lonchura cucullata* | 0.84 | *Accipiter superciliosus* | 0.71 |
| *Luscinia cyane* | 0.84 | *Aimophila rufescens* | 0.71 |
| *Megascops choliba* | 0.84 | *Amazonetta brasiliensis* | 0.71 |
| *Microcerculus bambla* | 0.84 | *Andropadus milanjensis* | 0.71 |
| *Mimus saturninus* | 0.84 | *Anthreptes fraseri* | 0.71 |
| *Momotus momota* | 0.84 | *Anthreptes platurus* | 0.71 |
| *Myiarchus ferox* | 0.84 | *Basileuterus rufifrons* | 0.71 |
| *Myiozetetes cayanensis* | 0.84 | *Bradornis microrhynchus* | 0.71 |
| *Myrmeciza atrothorax* | 0.84 | *Charadrius pecuarius* | 0.71 |
| *Myrmothera campanisona* | 0.84 | *Cisticola tinniens* | 0.71 |
| *Nectarinia senegalensis* | 0.84 | *Clamator levaillantii* | 0.71 |
| *Oporornis agilis* | 0.84 | *Erythrura trichroa* | 0.71 |
| *Oriolus chinensis* | 0.84 | *Eubucco versicolor* | 0.71 |
| *Perdix perdix* | 0.84 | *Florisuga mellivora* | 0.71 |
| *Petronia dentata* | 0.84 | *Hirundo rupestris* | 0.71 |
| *Ploceus nigerrimus* | 0.84 | *Hylophylax punctulatus* | 0.71 |
| *Prinia subflava* | 0.84 | *Hypocnemis flavescens* | 0.71 |
| *Psarocolius viridis* | 0.84 | *Leucopternis kuhli* | 0.71 |
| *Struthidea cinerea* | 0.84 | *Lonchura castaneothorax* | 0.71 |
| *Tockus nasutus* | 0.84 | *Meliphaga aruensis* | 0.71 |
| *Trachyphonus vaillantii* | 0.84 | *Myzomela obscura* | 0.71 |
| *Vireo solitarius* | 0.84 | *Nectarinia amethystina* | 0.71 |
| *Accipiter nisus* | 0.83 | *Oriolus oriolus* | 0.71 |
| *Ailuroedus buccoides* | 0.83 | *Parus ater* | 0.71 |
| *Aimophila ruficeps* | 0.83 | *Pogoniulus scolopaceus* | 0.71 |
| *Alectoris chukar* | 0.83 | *Pteroglossus aracari* | 0.71 |
| *Anisognathus somptuosus* | 0.83 | *Ramphastos dicolorus* | 0.71 |
| *Atlapetes pileatus* | 0.83 | *Sitta pygmaea* | 0.71 |
| *Catharus dryas* | 0.83 | *Sylvia deserticola* | 0.71 |
| *Cracticus torquatus* | 0.83 | *Taeniopygia guttata* | 0.71 |
| *Criniger calurus* | 0.83 | *Tangara cyanoptera* | 0.71 |
| *Dicrurus adsimilis* | 0.83 | *Thryothorus longirostris* | 0.71 |
| *Euphonia pectoralis* | 0.83 | *Trogon rufus* | 0.71 |
| *Euplectes hartlaubi* | 0.83 | *Turdus fuscater* | 0.71 |
| *Francolinus afer* | 0.83 | *Turdus olivater* | 0.71 |
| *Galerida cristata* | 0.83 | *Anthus lineiventris* | 0.7 |

(Continued) Bird species with high susceptibility to malaria predicted by interpretable machine learning models

| Species | Susceptibility | Species | Susceptibility |
|---|---|---|---|
| *Gymnopithys rufigula* | 0.83 | *Archilochus alexandri* | 0.7 |
| *Hippolais opaca* | 0.83 | *Attila phoenicurus* | 0.7 |
| *Hypocnemoides maculicauda* | 0.83 | *Campephilus melanoleucos* | 0.7 |
| *Lagonosticta rubricata* | 0.83 | *Campylopterus largipennis* | 0.7 |
| *Lonchura bicolor* | 0.83 | *Carpodacus pulcherrimus* | 0.7 |
| *Melierax gabar* | 0.83 | *Chrysuronia oenone* | 0.7 |
| *Motacilla alba* | 0.83 | *Cisticola guinea* | 0.7 |
| *Myiozetetes luteiventris* | 0.83 | *Cisticola rufilatus* | 0.7 |
| *Periporphyrus erythromelas* | 0.83 | *Eminia lepida* | 0.7 |
| *Pomatostomus isidorei* | 0.83 | *Estrilda atricapilla* | 0.7 |
| *Poospiza cabanisi* | 0.83 | *Falco naumanni* | 0.7 |
| *Pseudoscops clamator* | 0.83 | *Gymnopithys leucaspis* | 0.7 |
| *Pseudoseisura lophotes* | 0.83 | *Halcyon chelicuti* | 0.7 |
| *Pytilia afra* | 0.83 | *Jynx torquilla* | 0.7 |
| *Serinus burtoni* | 0.83 | *Laniarius ferrugineus* | 0.7 |
| *Sporophila americana* | 0.83 | *Lepidothrix iris* | 0.7 |
| *Sporophila caerulescens* | 0.83 | *Melanerpes cactorum* | 0.7 |
| *Zosterops montanus* | 0.83 | *Neocossyphus poensis* | 0.7 |
| *Accipiter tachiro* | 0.82 | *Phoenicircus carnifex* | 0.7 |
| *Asio stygius* | 0.82 | *Procnias nudicollis* | 0.7 |
| *Carduelis flavirostris* | 0.82 | *Rhegmatorhina hoffmannsi* | 0.7 |
| *Caryothraustes canadensis* | 0.82 | *Selasphorus platycercus* | 0.7 |
| *Catherpes mexicanus* | 0.82 | *Sporophila collaris* | 0.7 |
| *Cichladusa arquata* | 0.82 | *Tangara gyrola* | 0.7 |
| *Clamator jacobinus* | 0.82 | *Tapera naevia* | 0.7 |
| *Criniger barbatus* | 0.82 | *Toxostoma longirostre* | 0.7 |
| *Dendroica fusca* | 0.82 | *Vermivora ruficapilla* | 0.7 |
